# A redesigned TetR-aptamer system to control gene expression in *Plasmodium falciparum*

**DOI:** 10.1101/2020.05.19.105411

**Authors:** Krithika Rajaram, Hans B. Liu, Sean T. Prigge

## Abstract

One of the most powerful approaches to understanding gene function involves turning genes on and off at will and measuring the impact at the cellular or organismal level. This particularly applies to the cohort of essential genes where traditional gene knockouts are inviable. In *Plasmodium falciparum*, conditional control of gene expression has been achieved by using multi-component systems in which individual modules interact with each other to regulate DNA recombination, transcription or posttranscriptional processes. The recently devised TetR-DOZI aptamer system relies on the ligand-regulatable interaction of a protein module with synthetic RNA aptamers to control the translation of a target gene. This technique has been successfully employed to study essential genes in *P. falciparum* and involves the insertion of several aptamer copies into their 3’ untranslated regions (UTRs) which provide control over mRNA fate. However, aptamer repeats are prone to recombination and one or more copies can be lost from the system, resulting in a loss of control over target gene expression. We rectified this issue by redesigning the aptamer array to minimize recombination while preserving the control elements. As proof of concept, we compared the original and modified arrays for their ability to knock down the levels of a putative essential apicoplast protein (PF3D7_0815700) and demonstrated that the modified array is highly stable and efficient. This redesign will enhance the utility of a tool that is quickly becoming a favored strategy for genetic studies in *P. falciparum*.

**Importance:** Malaria elimination efforts have been repeatedly hindered by the evolution and spread of multidrug-resistant strains of *Plasmodium falciparum*. The absence of a commercially available vaccine emphasizes the need for a better understanding of *Plasmodium* biology in order to further translational research. This has been partly facilitated by targeted gene deletion strategies for the functional analysis of parasite genes. However, genes that are essential for parasite replication in erythrocytes are refractory to such methods, and require conditional knockdown or knockout approaches to dissect their function. One such approach is the TetR-DOZI system that employs multiple synthetic aptamers in the untranslated regions of target genes to control their expression in a tetracycline-dependent manner. Maintaining modified parasites with intact aptamer copies has been challenging since these repeats are frequently lost by recombination. By interspacing the aptamers with unique sequences, we created a stable genetic system that remains effective at controlling target gene expression.

## Introduction

Significant strides have been made towards reducing the global burden of malaria and associated mortality since the turn of the millennium. However, progress towards eradication has slowed down in the last few years and could be further undermined by the emergence of resistant strains of *Plasmodium falciparum* to many frontline antimalarials (1). Understanding the genetic and mechanistic basis of resistance, along with the identification of novel drug targets, is therefore vital for the advancement of antimalarial therapies. These efforts have been aided by a considerable improvement in annotation of the *P. falciparum* genome since its release in 2002, with known or putative functions assigned to over 65% of predicted genes (2). Several technological advances in recent years have changed the landscape of molecular genetics in *P. falciparum* from relative intractability to the experimental validation of almost a quarter of the total number of protein-coding genes. Many of these studies have yielded unexpected results. For instance, several seemingly promising drug targets were found to be completely dispensable during the symptomatic blood stage of the parasite life cycle (3-5). There are also indications that many functionally annotated proteins have acquired additional or alternative roles in *Plasmodium* (6-11). These examples highlight a continuing need for the characterization of *Plasmodium* genes in their native context along with the improvement and expansion of our toolkit of genetic techniques.

Gene deletion studies have been greatly facilitated by the introduction of CRISPR/Cas9 to *P. falciparum*. However, knockouts are applicable only to the cohort of genes whose products are either dispensable during intraerythrocytic development or can be chemically or genetically bypassed (12-15). To address this, several conditional gene expression strategies have been developed that can be integrated with CRISPR/Cas9 for the functional analysis of essential blood-stage genes.

Nearly all conditional systems in *Plasmodium* involve the binding of small molecule ligands to nucleic acid/protein modules to control gene expression or protein function. Inducible FKBP-FRB dimerization by rapamycin and its analogs (rapalogs) forms the basis for two such techniques, 1) conditional DiCre recombinase-mediated excision of target genes that are flanked by loxP sites, and 2) conditional mislocalization of FKBP fusion proteins, known as knock sideways (KS) (16-18). Destabilizing FKBP (ddFKBP) or hDHFR (human dihydrofolate reductase) domains have been used to confer ligand-dependent stability to *Plasmodium* proteins (19-21). In the absence of ligand, these fusion proteins are degraded by the proteasome. N-terminal fusions of recently developed FKBP variants can traffic proteins to the lumen of the apicoplast organelle in a rapalog-controllable fashion (22). These tools vary in dynamic range and their successful application to a target gene may be influenced by gene expression levels, subcellular protein location or tolerance of the protein to N or C-terminal modifications.

To overcome constraints imposed by protein structure and location, methods have been developed to control gene expression at the post-transcriptional level in *P. falciparum*, either by inducible degradation or sequestration of mRNA. The self-cleaving GlmS ribozyme has been successfully appended to the 3’ UTR of target genes and the resulting transcripts are cleaved in a glucosamine-dependent manner (23). Various tetracycline repressor protein (TetR)-based systems have been tested in *P. falciparum*, but perhaps the most successful strategy involves the regulatable interaction of a TetR-DOZI fusion protein with a synthetic RNA aptamer array (24). TetR normally binds to its cognate dsDNA operator sequences to block the transcription of a target gene. This repression can be reversed by the allosteric interaction of tetracycline (TC) or its analogs with TetR. To adapt the TetR system to post-transcriptional regulation, Belmont and Niles (2010) conducted a screen for RNA aptamers that bind to TetR in a TC-regulatable fashion (25). Candidate aptamers that fulfilled these parameters were predicted to form stem-loop structures with conserved sequence motifs in the loops. Placing an RNA aptamer in the 5’ UTR of reporter genes resulted in a TC-dependent, 3 to 5-fold change in expression levels. To increase the dynamic range of the TetR-aptamer system, Ganesan et al. (2016) exploited the native translational regulation capabilities of *Plasmodium* (24). DOZI (‘development of zygote inhibited’) is a *Plasmodium* RNA helicase that represses translation by assembling specific transcripts into inactive messenger ribonucleoprotein particles (mRNPs) (26). The integration of DOZI with the TetR-aptamers afforded much greater control over reporter gene expression, especially when the 5’ UTR aptamer was combined with multiple aptamer copies in the 3’ UTR (20 to 300-fold change in induction) (24).

Since it can be challenging to simultaneously modify the 5’ and 3’ UTRs of endogenous genes, several groups have effectively utilized TetR-DOZI with an aptamer array inserted only into the 3’ UTR (5, 24, 27-42). The 3’ UTR array consists of 10 aptamer copies, separated by spacers that are identical in sequence and length. Such tandem repeats are inherently unstable and tend to be deleted by recombination. We and others have observed truncations in the aptamer array with a consequent loss of control over gene expression, thereby limiting the utility of this knockdown strategy (28, 29, 34, 35). To rectify this, we modified the array to contain unique spacers while retaining the secondary structure of the aptamers. We validated the redesigned aptamer array by targeting a putative apicoplast ubiquitin-like protein (PF3D7_0815700), referred to as PfAUBL. We found that the new array does not lose its aptamers during plasmid propagation in *E. coli* or upon insertion into the *P. falciparum* genome. The redesign does not interfere with aptamer function, as demonstrated by efficient TC-dependent knockdown of the PfAUBL protein that subsequently results in parasite death, indicating an essential role for PfAUBL. The enhanced stability of the aptamer array will expand the usage of TetR-DOZI for gene function studies in *P. falciparum*.

## Results

### Aptamer arrays undergo truncation in *E. coli* and in *P. falciparum*

The TetR-DOZI aptamer system has been successfully used for the conditional knockdown of several *P. falciparum* proteins. We implemented this approach to study a putative apicoplast protein called PfAUBL (PF3D7_0815700). PfAUBL was identified as essential for blood-stage survival in the genome-wide *piggyBac* mutagenesis screen conducted in *P. falciparum* (43). The protein contains an N-terminal apicoplast transit peptide and a ubiquitin-like domain (44). We utilized a previously published plasmid pMG74 for the insertion of a 10X aptamer array downstream of the PfAUBL coding region (24). Plasmids from individual bacterial colonies were analyzed by PCR to verify the size of the aptamer array. While some plasmids contained a full-length (≈1 kb) array, other plasmids appeared to have lost a few aptamers from their arrays (**Fig. 1A**). We selected a plasmid with an intact sequence-confirmed array to generate PfAUBL^Apt^ parasites. Correct insertion of the aptamer array at the 3’ UTR of PfAUBL^Apt^ was confirmed by PCR (**Fig. 1B, Fig. S1 and Fig. S2A**).

**Figure 1.**
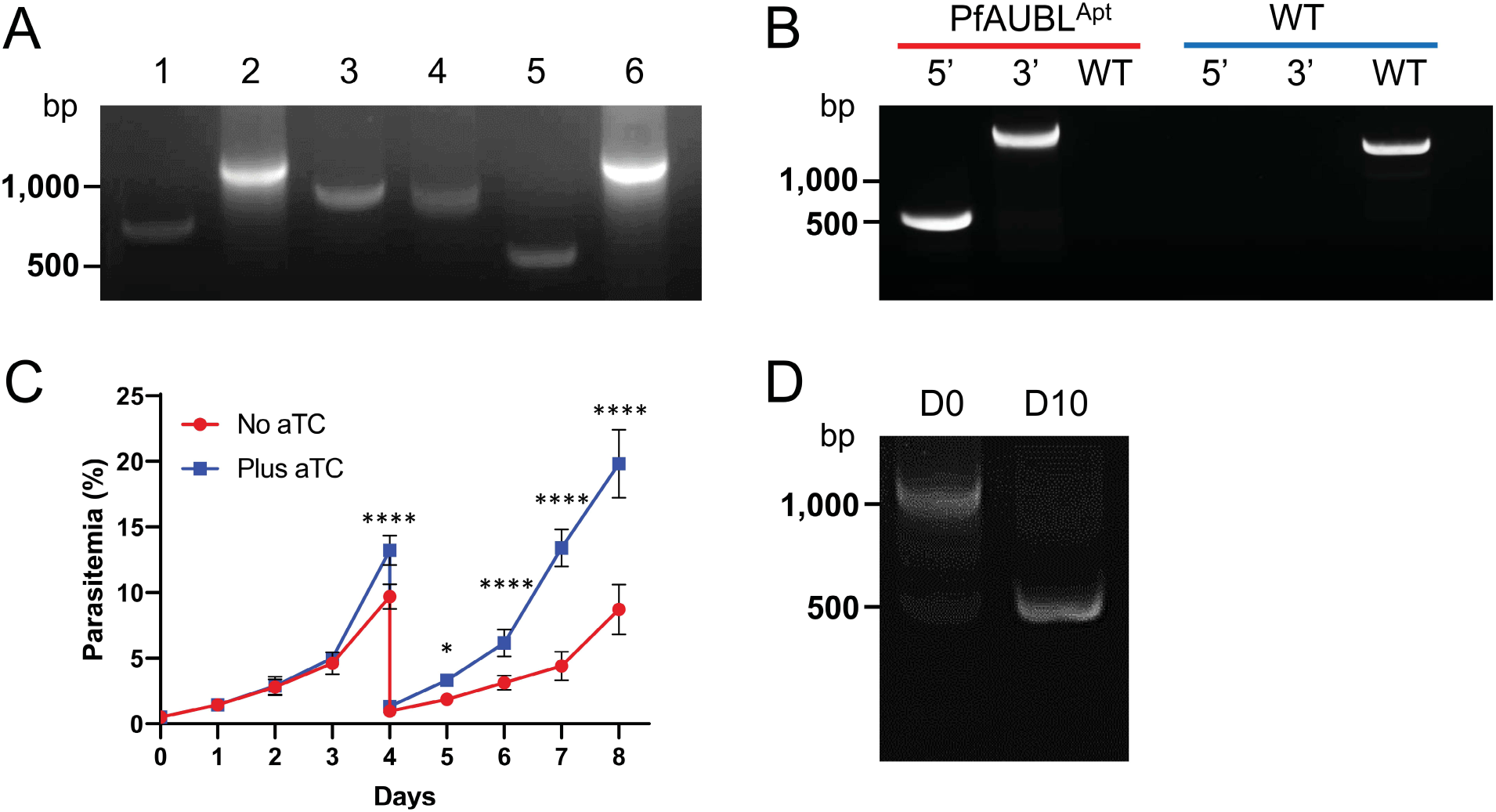
Aptamer arrays are unstable during propagation in *E. coli* and *P. falciparum*. A) PCR amplification of the aptamer array from six bacterial colonies transformed with pMG74^PfAUBL^ plasmid. PCR products of full-length arrays are ≈1 kb in size. Smaller products in some lanes signify truncated arrays. B) Integration of pMG74^PfAUBL^ plasmid into the 3’ UTR of the PfAUBL gene was confirmed by PCR amplification of the 5’ and 3’ loci. PCR for the unmodified locus failed to detect any residual wild-type parasites (red) in the PfAUBL^Apt^ population; wild-type parasites served as a control (blue) C) Growth of PfAUBL^Apt^ parasites in the presence or absence of aTC was monitored by daily blood smears for 8 days. The parasites were cut 1:10 on day 4. Removal of aTC caused moderate growth inhibition of PfAUBL^Apt^ parasites beginning on day 4 (two-way ANOVA, followed by Bonferroni’s correction; ****, *P* < 0.0001; **, *P* < 0.01; *, *P* < 0.05). Error bars represent the standard error of the mean from two independent experiments, each conducted in quadruplicate. D) PCR amplification of the aptamer array from the aTC-treated PfAUBL^Apt^ population shows 1 kb (full-length) and 0.5 kb products, indicating that some parasites have lost a few aptamers from their array (D0). Surviving parasites at 10 days post aTC removal (D10) have a predominant 0.5 kb band, consistent with a truncated array.

Polyclonal parasite populations from two independent transfections were tested for growth in the presence and absence of anhydrotetracycline (aTC), a tetracycline analog. Both parasite lines exhibited moderate growth inhibition 4-5 days after the removal of aTC, but continued to multiply over the course of the growth assay (**Fig. 1C** and **Fig. S2B**). To determine if the parasites replicating without aTC still contained all 10 aptamers, we performed PCR to determine the size of the array before and after aTC removal. An expected ≈1 kb band was the predominant product from aTC-treated parasites (**Fig. 1D and Fig. S2C**). However, one or more smaller bands were also observed, which indicated that both transfections yielded a mixed population of parasites containing varying sizes of aptamer arrays. Post aTC removal, only the smaller PCR products corresponding to truncated arrays could be amplified from the surviving parasites. This suggests that PfAUBL is an essential protein and that parasites with a fully functional aptamer array died upon withdrawal of aTC. Presumably, parasites with arrays that contained fewer aptamers produced enough PfAUBL to survive in the absence of aTC.

The extended time interval between parasite transfection and phenotypic analysis of transgenic lines could enable the gradual accumulation of parasites with recombined aptamer arrays. We therefore attempted to isolate a clonal parasite population with an intact array from one of the transfections. PCR analysis of five putative clones revealed aptamer arrays of varying sizes. Parasites from well E11 were found to be nonclonal, and none of the remaining samples contained full-length arrays (**Fig. 2A**). Parasites from clone E10 had the largest array with 8 aptamers and were therefore analyzed further. Clone E10 parasites grown in the absence of aTC exhibited severe growth inhibition beginning on Day 5, and this defect was much more pronounced than the previously analyzed mixed populations (**Fig. 2B**). However, a small proportion of aTC-depleted parasites continued to grow and replicate. When examined by PCR on Day 10, the surviving parasites were found to contain a truncated array with 5 aptamers, which was confirmed by sequencing (**Fig. 2C**). This indicates that aptamer loss can occur within a short time frame and confers a selective advantage to parasites in non-permissive growth conditions.

**Figure 2.**
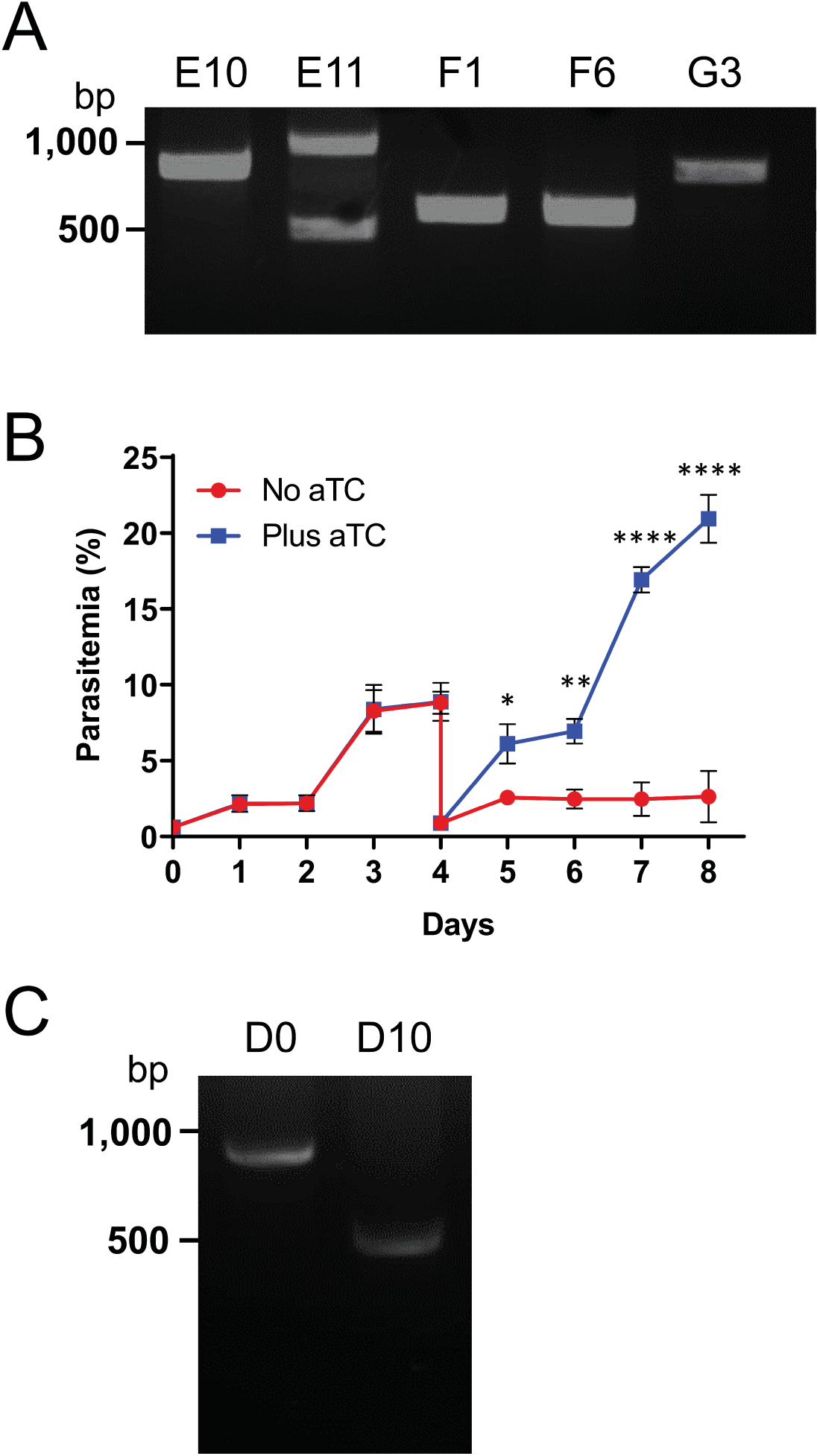
Aptamers of a PfAUBLApt clone are rapidly lost under non-permissive growth conditions. A) PCR amplification of aptamer arrays from cloned parasite populations reveals varying array sizes. B) Growth of Clone E10 parasites in the presence or absence of aTC was monitored by flow cytometry for 8 days. The parasites were cut 1:10 on day 4. Removal of aTC caused significant growth inhibition of parasites beginning on day 5 (two-way ANOVA, followed by Bonferroni’s correction; ****, *P* < 0.0001; **, *P* < 0.01; *, *P* < 0.05). Error bars represent the standard error of the mean from two independent experiments, each conducted in quadruplicate. C) PCR amplification of the aptamer array from aTC-treated Clone E10 parasites (D0) shows the expected 0.8 kb product, corresponding to 8 aptamers. Parasites surviving after 10 days of no aTC treatment (D10) have a predominant 0.5 kb band, indicating that more aptamers have been lost from the original 8-aptamer array found in Clone E10.

### Modification of spacers between RNA aptamers

The TetR-aptamer array contains 10 copies of an aptamer separated by identical spacers (86 bp each and 860 bp in total). Such repetitive sequences are prone to recombination events that can lead to truncations. To minimize recombination between individual aptamers and spacers, we made the following modifications to the array. The original aptamers are 51 bp in length and are predicted to fold into stem-loop structures (**Fig. 3A**). Bases 1-10 and 42-51 are palindromic and can pair with each other to form a stem with a predicted melting temperature of ≈33°C. Bases 1-6 and 45-51 also encode BamHI restriction enzyme sites (**Fig. 3A**). To facilitate future cloning reactions that may require BamHI, we altered these bases while maintaining the same GC content (and therefore a similar melting temperature). Each modified aptamer now contained a unique palindromic 6 bp sequence on either end. The loop regions containing conserved sequence motifs that are known to be important for TetR binding were left unaltered. The aptamers in the original array were separated by identical 35 bp spacers. We reduced the spacer length to 11 bp and altered the sequence such that each spacer was unique and formed no significant secondary structure with the 10 stem-loop aptamers (**Fig. 3B**). Thus, each of the array elements now contains 23 bp of unique sequence corresponding to the spacer region (11 bp) and part of the double stranded stem (12 bp). The redesigned spacers have higher GC content than the original spacer (52% versus 44%), thus facilitating the design of unique primers for sequencing or for future genetic manipulations. They also do not contain any common restriction endonuclease sites that could interfere with downstream cloning. We replaced the original 860 bp array in the pMG74 plasmid with the redesigned 619 bp array (**Fig. 3C**) to generate the pKD vector.

**Figure 3.**
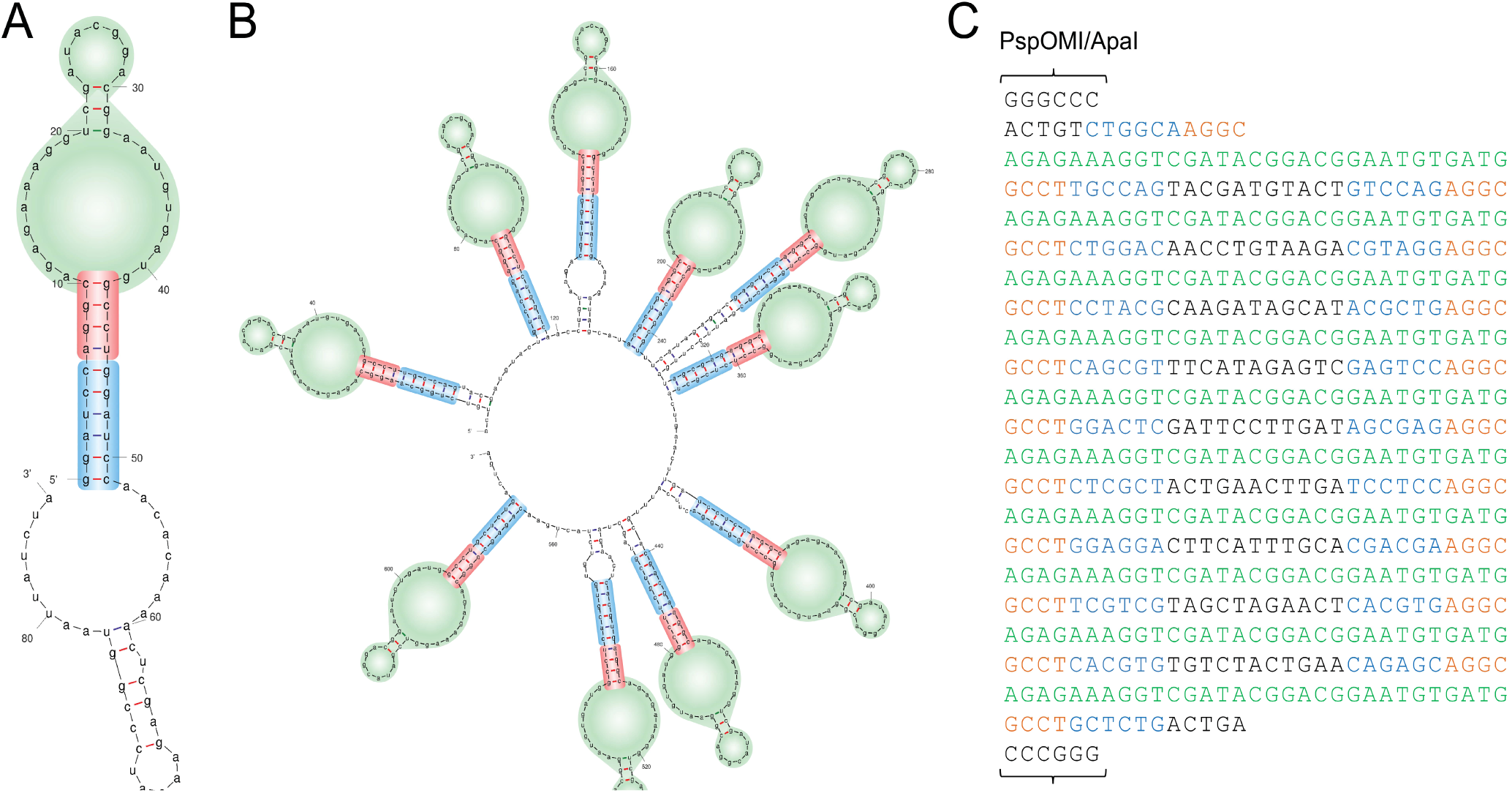
Spacers in a redesigned RNA aptamer array do not have significant secondary structure. A) Secondary structure prediction of a single RNA aptamer from the original array using mFold. The 51 bp stem-loop structure contains a loop responsible for TetR binding (green) and a double stranded stem (red), in part formed from residues corresponds to a BamHI site (blue). The stem-loop structure is followed by a 35 bp spacer. B) Redesigned aptamer array which preserves the ten stem-loop structures, but separates each aptamer with unique 11 bp spacer sequences. The BamHI sequences found in the original aptamers were replaced with unique 6 bp sequences with the same GC content (blue). C) Sequence of the new array flanked by endonuclease sites with color coding corresponding to that used in B). Each of the array elements now contains 23 bp of unique sequence corresponding to the spacer region (black) and part of the double stranded stem (blue).

### The redesigned aptamer array exhibits enhanced stability in *E. coli* and *P. falciparum*

pKD plasmids with PfAUBL homology arms were isolated from a few bacterial colonies and their aptamer arrays were examined by PCR. All tested plasmids were found to contain an intact array (≈0.7 kb) (**Fig. 4A**). Parasites were transfected as before with a sequence-confirmed pKD plasmid to generate PfAUBL^NewApt^ parasites. The insertion of the new aptamer array at the 3’UTR of PfAUBL was confirmed by PCR (**Fig. 4B and Fig. S1**). To determine if the aptamers in the redesigned array retained responsiveness to aTC, PfAUBL^NewApt^ parasites were subjected to a growth assay in the presence or absence of aTC. Growth was completely abrogated 4 days after the withdrawal of aTC, and parasites were undetectable in cultures beyond day 5 (**Fig. 4C**). The flow cytometric results were confirmed by manual inspection of blood smears. PCR analysis of parasites 4 days post aTC removal did not indicate any loss of aptamers from the array (**Fig. 4D**). Cultures depleted of aTC were maintained for 21 additional days, but no parasite recrudescence was observed. These results implied that the use of unique spacers between identical aptamers was sufficient to prevent recombination within the array and that the changes that we made to the aptamer array did not affect the functionality of the knockdown system. PfAUBL^NewApt^ parasites maintained in permissive culture conditions for several months did not exhibit any loss of aptamers.

**Figure 4.**
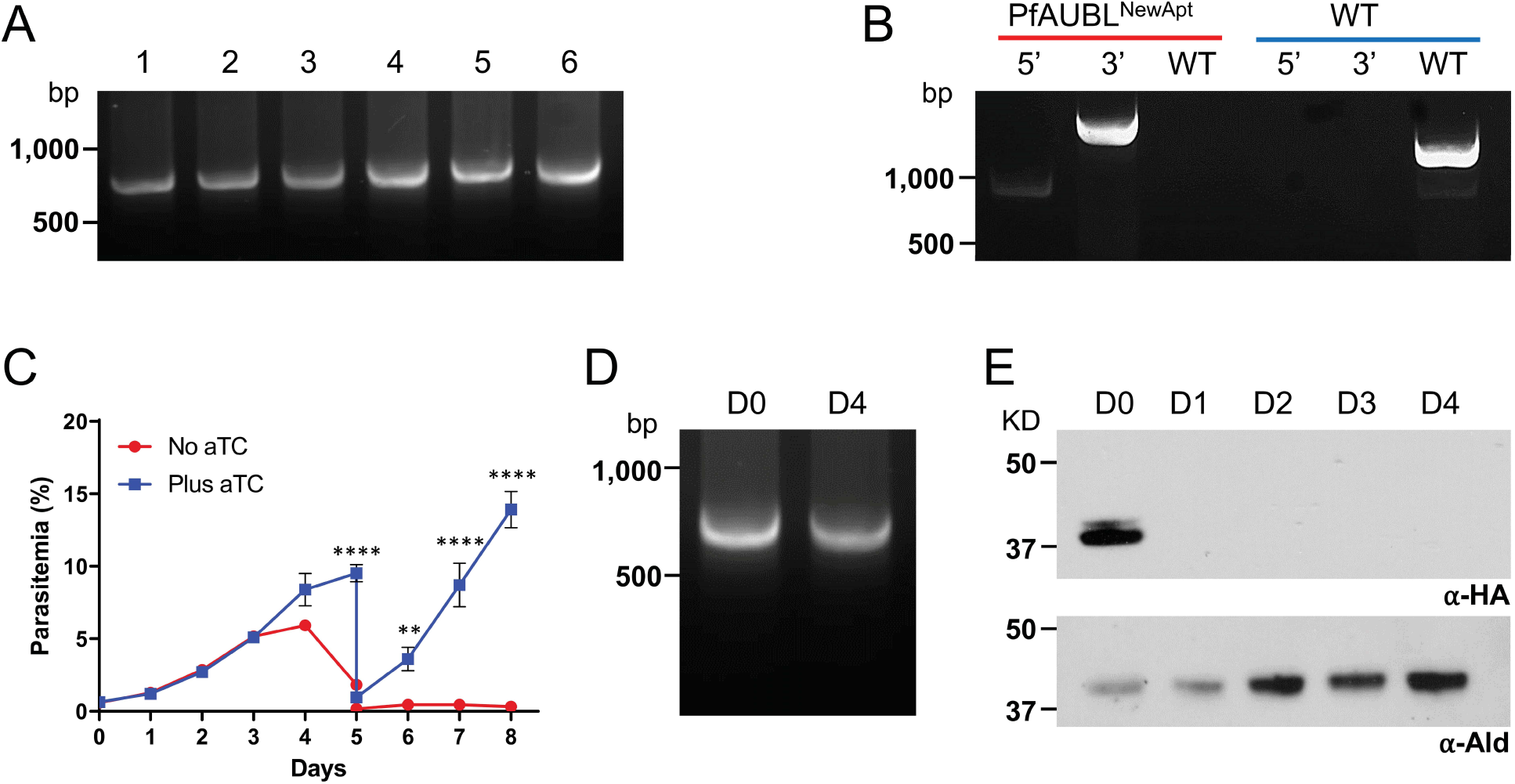
The redesigned aptamer array is stably maintained and retains responsiveness to aTC. A) PCR amplification of the aptamer array from six bacterial colonies transformed with pKD^PfAUBL^ plasmid shows that all colonies contain full-length (≈0.7 kb) arrays. B) Integration of pKD^PfAUBL^ plasmid into the 3’ UTR of the PfAUBL gene was confirmed by PCR amplification of the 5’ and 3’ loci. PCR for the unmodified locus failed to detect any residual wild-type parasites (red) in the PfAUBL^NewApt^ population; wild-type parasites served as a control (blue) C) Growth of PfAUBL^NewApt^ parasites in the presence or absence of aTC was monitored by flow cytometry for 8 days. The parasites were cut 1:10 on day 5. Removal of aTC caused significant growth inhibition of PfAUBL^NewApt^ parasites beginning on day 5 (two-way ANOVA, followed by Bonferroni’s correction; ****, *P* < 0.0001; **, *P* < 0.01; *, *P* < 0.05). Error bars represent the standard error of the mean from two independent experiments, each conducted in quadruplicate. D) PCR amplification of the aptamer array from aTC-treated PfAUBL^NewApt^ parasites resulted in a single band of 0.7 kb corresponding to the full-length array (D0). DNA from dying or dead parasites at 4 days post aTC removal (D4) also generated a 0.7 kb PCR product. E) Western blotting using anti-HA antibody to probe for PfAUBL protein shows a rapid decrease in protein levels upon aTC removal. Anti-aldolase antibody was used as a loading control.

The pMG74 plasmid was designed to append 1X HA and 1X FLAG tags onto the C-terminus of the target protein. However, we failed to detect PfAUBL in PfAUBL^Apt^ parasites using anti-HA or anti-FLAG antibodies. To enhance detection capabilities, we used a 3X HA tag in the pKD plasmid. Probing for PfAUBL with anti-HA antibody demonstrated robust protein detection in aTC-treated PfAUBL^NewApt^ parasites (**Fig. 4E**). To determine the efficiency of protein knockdown, we examined the levels of PfAUBL in aTC-depleted parasites over a 4-day period. Protein levels were undetectable 24 hours post aTC removal, indicating that the knockdown was quick and efficient.

## Discussion

Recent years have witnessed a steady development of genetic techniques for the functional analysis of essential genes in *P. falciparum*. The TetR-DOZI aptamer tool for conditional gene knockdown is particularly promising because it is unhindered by structural properties or the subcellular localization of target proteins. It can also be used to control gene expression in a reversible and tunable manner by varying the dosage of the tetracycline ligand, thus permitting the evaluation of null as well as hypomorphic phenotypes. However, its broad implementation has been limited by two key issues. One is that large dynamic range control of gene expression is achieved only when aptamers are introduced on either side of the target gene’s coding region (24). While this is easy to accomplish with transgenes, modifying both the 5’ and 3’ UTRs of native genes is less straightforward. The use of a 10X aptamer array in the 3’ UTR alone has been sufficient to knock down the expression of several genes, but it may not be enough to achieve complete conditional control of genes that are expressed at high levels. A recent publication overcomes this issue by utilizing a linear vector that carries a recodonized copy of the target gene and homology arms corresponding to the 5’ and 3’ UTRs, thereby allowing simultaneous insertion of aptamers into the upstream and downstream sequences (42).

This study addresses a second drawback of the TetR-DOZI system, namely the instability of the 3’ UTR aptamer array due to its repetitive nature. Our results show that truncated arrays arise frequently and that arrays below a certain threshold of aptamers are unable to exert sufficient control over gene expression. This is supported by the original study describing the TetR-RNA aptamers in which a parallel analysis of arrays containing 5 or 10 aptamers showed that the 5X array was unable to regulate reporter gene expression in yeast (24). By replacing the identical spacers in between aptamers with unique sequences, we were able to avoid array truncations. Importantly, modification of the spacers did not interfere with the functionality of the aptamers. We demonstrated this by successfully knocking down the expression of the ubiquitin-like protein PfAUBL, which led to parasite death. The redesigned aptamer array can also be employed in the linear vector described above to facilitate the installation of aptamers in both the 5’ and 3’ UTRs. The combination of these techniques will greatly expand the application of the TetR-DOZI system in efforts to assign functions to the hundreds of uncharacterized essential *P. falciparum* genes.

## Methods

### Parasite culture

*P. falciparum* Nf54^attB^ parasites (45) were cultured in human erythrocytes at 2% hematocrit in CMA (Complete Medium with Albumax) containing RPMI 1640 medium with L-glutamine (USBiological Life Sciences), supplemented with 20 mM HEPES, 0.2% sodium bicarbonate, 12.5 µg/mL hypoxanthine, 5 g/L Albumax II (Life Technologies) and 25 µg/mL gentamicin. Cultures were maintained at 37°C in a gas mixture containing 94% N_2_, 3% O_2_ and 3% CO_2_.

### Plasmid construction

The pMG74 plasmid was digested with AatII and XmaI to remove 1X HA and 1X FLAG tags and the existing aptamer array (24). A small synthetic DNA fragment containing cloning sites for the aptamer array (PspOMI and XmaI) and residues encoding a 3X HA tag was inserted into pMG74 using the AatII and XmaI sites (LifeSCT). The redesigned 619 bp aptamer array (**Fig. 3C**) was cut with PspOMI and XmaI from a synthetic plasmid (LifeSCT) and ligated into the same sites in pMG74. The resulting plasmid was called pKD and contains the homology arm cloning sites, 3X HA tag and aptamer array shown in **Fig. S3A**.

To create PfAUBL knockdown constructs, pMG74 and pKD were digested with AscI and AatII. Homology arms HA1 and HA2 of the PfAUBL gene were amplified from *P. falciparum* genomic DNA using the primer pairs AUBL HA1.kd-F + AUBL HA1.kd-R and AUBL HA2.kd-F + AUBL HA2.kd-R, respectively (**Table S1**). The homology arms were fused in an additional PCR reaction using the primer pair AUBL HA2.kd-F + AUBL HA1.kd-R, creating a combined HA2-HA1 fragment separated by two EcoRV sites. This fragment was inserted into digested pMG74 and pKD plasmids to generate pMG74^PfAUBL^ and pKD^PfAUBL^, respectively. Prior to transfection, these plasmids were linearized with EcoRV.

The Cas9 enzyme and the guide RNA were expressed from a modified version of pUF1-Cas9 (46). Plasmid pUF1-Cas9 (11,096 bp) was modified in several steps to accommodate the insertion of the guide RNA expression cassette found in pRS-LacZ (47). In the first step, the promoter driving Cas9 expression (Hsp86) was removed using XhoI and AflII and replaced with an adaptamer formed from 5’-phosphorylated primers pCas.XhoBtg.F and pCas.XhoBtg.R (**Table S1**). This plasmid was then digested with XhoI/BtgZI, blunted with T4 polymerase and ligated. The resulting plasmid is smaller (10,221bp) and brings Cas9 expression under the control of the bidirectional calmodulin promoter. In the second step, a 340 bp region of the bluescript plasmid backbone containing several endonuclease sites was removed through digestion with EcoRI/AatII, blunted with T4 polymerase and ligated. In the third step, digestion with XmaI/SapI was used to remove the 3’ UTR of Cas9 (PbDHFR-TS) and a complete lac operon found in the bluescript plasmid backbone. This region was replaced with an adaptamer formed from the 5’-phosphorylated primers pUF.NotI.F and pUF.NotI.R (**Table S1**) containing a NotI site for insertion of the guide RNA cassette found in pRS-LacZ. Before this insertion could take place, however, two BsaI sites in the plasmid backbone had to be removed (BsaI will be used later to insert guide RNA sequences). The plasmid was digested with BsaI, removing a 1,992 bp region containing both recognition sites for this Type IIs endonuclease. PCR primers BsaMut.GG.F and BsaMut.GG.R (**Table S1**) were designed with flanking BsaI sites for Golden Gate insertion back into the plasmid, but with two single base changes to alter the original two BsaI recognition sites. Thus, after digestion of the PCR product with BsaI, and ligation back into the cut plasmid, the resulting plasmid was identical in size (8,757 bp) and contained two single nucleotide changes that removed both BsaI recognition sites. The guide RNA expression cassette from plasmid pAIO (29) was then excised with NotI and inserted into the same site bidirectionally, generating a forward and a reverse orientation. The orientation with the U6 5’ regulatory element adjacent to Cas9 was chosen with the expectation that this region would act as a 3’ UTR for Cas9 (it contains the 3’ UTR of the ribosomal L18 protein, PF3D7_1341200). Finally, the plasmid was digested with BtgZI to remove the pre-guide RNA and the LacZ expression cassette from pRS-LacZ (47) was excised with BsaI and inserted into this site. The final pCasG-LacZ plasmid (10,886) is smaller than the original pUF1-Cas9 plasmid and contains a guide RNA expression cassette that can be used for blue/white colony screening after insertion of a guide RNA sequence with BsaI (**Fig. S3B**).

To generate a pCasG-LacZ plasmid with a guide RNA sequence that targets PfAUBL, pCasG-LacZ was digested with BsaI. PfAUBL-specific guide RNAs were synthesized as oligos (**Table S1**), annealed, and inserted into the digested plasmid using In-Fusion.

### Parasite transfections

*P. falciparum* NF54^attB^ parasites were transfected using a previously described method with minor modifications (48). Briefly, 350 µL of red blood cells were electroporated with 75 µg each of the pCasG^PfAUBL^ plasmid and either the linearized pMG74^PfAUBL^ or pKD^PfAUBL^ plasmids. The electroporated RBCs were mixed with 1.5 mL of a mixed-stage culture at ≈4% parasitemia. Transfected cultures were maintained in CMA with 0.5 µM anhydrotetracycline (aTC) (Cayman Chemical) for the first 48 hours, after which they were cultured in selective media containing 2.5 µg/mL Blasticidin (Corning) and 1.5 µM DSM-1 (Calbiochem) along with aTC for 7 days. Cultures were then switched to CMA with aTC until the reemergence of parasites, upon which they were transferred to CMA containing Blasticidin and aTC. Single clones of PfAUBL^Apt^ parasites were isolated by limiting dilution in a 96-well plate.

### Genotyping and aptamer PCR

A 3 µL volume of parasite culture or 10 ng of plasmid was used in 25 µL PCR reactions with Phusion High-Fidelity DNA polymerase (NEB). To confirm pMG74^PfAUBL^ insertion into the 3’ UTR of PfAUBL, the following primer pairs were used: AUBL 5’ F + pMG74 R for the 5’ integration locus, and TetR-seq + AUBL 3’ R for the 3’ integration locus. To confirm pKD^PfAUBL^ insertion into the 3’ UTR of PfAUBL, the following primer pairs were used: AUBL 5’ F + NewApt-5R for the 5’ integration locus, and TetR-seq + AUBL 3’ R for the 3’ integration locus. To determine if any wild-type parasites were still present in the transfected populations, the primer pair AUBL 5’ F + AUBL WT 3’R was used.

The original aptamer array in the pMG74 plasmid and PfAUBL^Apt^ parasites was amplified using the primer pair Apt-1F + Apt-10R. The redesigned aptamer array in the pKD plasmid and PfAUBL^NewApt^ parasites was amplified using the primer pair NewApt-1F + NewApt-10R.

### Growth assays

Parasites were washed with 10 mL CMA three times to remove aTC from the culture medium and then cultured in CMA alone or CMA with 0.5 µM aTC. They were seeded in a 96-well plate (Corning) at a 0.5% starting parasitemia and 2% hematocrit in a total volume of 250 µL per well, in quadruplicate for each medium condition. The plates were incubated in chambers gassed with 94% N_2_, 3% O_2_, 3% CO_2_ at 37°C. A 10 µL volume of each parasite sample was collected every day for blood smear analysis or flow cytometry, and the culture medium was exchanged every other day for eight days.

For growth curve determination by flow cytometry, samples collected on days 0-4 were diluted 1:10 in PBS and stored in a 96-well plate at 4°C. On day 4, the parasites were stained with SYBR Green by transferring 10 µL of the 1:10 dilutions to a 96-well plate containing 100 µL of 1X SYBR Green (Invitrogen) per well in PBS. The plates were incubated at room temperature under gentle shaking in the dark for 30 minutes. Post-incubation, 150 µL of PBS was added to each well to dilute unbound SYBR Green dye. The samples were analyzed with an Attune Nxt Flow Cytometer (Thermo Fisher Scientific), with a 50 µL acquisition volume, and a running speed of 25 µL/minute with 10,000 total events collected. The process was repeated on Day 8 for samples collected on days 5-8.

### Western blotting

Parasites were washed with CMA three times to remove aTC from the culture medium. Samples were collected immediately post wash, and then every 24 hours for the next 4 days. The collected samples were centrifuged at 500g for 5 min at room temperature (RT) and pellets were stored at −20°C until the time of protein extraction. To isolate parasites from RBCs, the pellets were thawed and resuspended in 0.15% saponin for 5 min at RT. Lysed RBCs were removed by washing three times with 1X PBS. Parasite pellets were resuspended in NuPAGE LDS sample buffer (Thermo Fisher) containing 2% ß-mercaptoethanol and boiled for 5 min. Proteins were resolved by SDS-PAGE on 4-12% gradient gels and transferred to nitrocellulose membranes. Membranes were blocked in 5% milk and probed overnight at 4°C with 1:2,500 rat anti-HA mAb 3F10 (Roche). They were then incubated for an hour at RT with 1:5,000 anti-rat horseradish peroxidase conjugated antibody (GE Healthcare). Protein bands were detected on X-ray film using SuperSignal West Pico Chemiluminescent Substrate (Thermo Scientific), according to the manufacturer’s protocol. For loading controls, membranes were stripped of antibody with 200 mM glycine (pH 2.0) for 5 min and reprobed with 1:25,000 mouse anti-aldolase mAB 2E11. After incubation with 1:10,000 anti-mouse horseradish peroxidase conjugated antibody (GE Healthcare), proteins were detected as described above.

## Acknowledgements

We thank David Fidock (Columbia University) for providing the NF54*att*B strain of *P. falciparum* parasites, Jaquin Niles (Massachusetts Institute of Technology) for the pMG74 plasmid, Josh Beck (Iowa State University) for the pAIO plasmid, and David Sullivan (Johns Hopkins Bloomberg School of Public Health) for mouse anti-aldolase mAB 2E11. This work was supported by the National Institutes of Health R01 AI065853 (STP), the Johns Hopkins Malaria Research Institute, and the Bloomberg Family Foundation. This work was also made possible by UL1 RR025005 from the NIH National Center for Research Resources. KR was supported by the NIH training grant T32AI007417.

## References

1. Anonymous. World malaria report 2019. World Health Organization.

2. Böhme U, Otto T, Sanders M, Newbold C, Berriman M. 2019. Progression of the canonical reference malaria parasite genome from 2002-2019 [version 2; peer review: 3 approved]. Wellcome Open Research 4.

3. Ke H, Sigala PA, Miura K, Morrisey JM, Mather MW, Crowley JR, Henderson JP, Goldberg DE, Long CA, Vaidya AB. 2014. The heme biosynthesis pathway is essential for *Plasmodium falciparum* development in mosquito stage but not in blood stages. The Journal of Biological Chemistry 289:34827–34837.

4. Shears MJ, Botté CY, McFadden GI. 2015. Fatty acid metabolism in the *Plasmodium* apicoplast: Drugs, doubts and knockouts. Molecular and Biochemical Parasitology 199:34–50.

5. Ke H, Ganesan SM, Dass S, Morrisey JM, Pou S, Nilsen A, Riscoe MK, Mather MW, Vaidya AB. 2019. Mitochondrial type II NADH dehydrogenase of *Plasmodium falciparum* (PfNDH2) is dispensable in the asexual blood stages. PLoS One 14:e0214023.

6. Jewett TJ, Sibley LD. 2003. Aldolase forms a bridge between cell surface adhesins and the actin cytoskeleton in apicomplexan parasites. Molecular Cell 11:885–894.

7. Buscaglia CA, Coppens I, Hol WGJ, Nussenzweig V. 2003. Sites of interaction between aldolase and thrombospondin-related anonymous protein in *Plasmodium*. Molecular Biology of the Cell 14:4947–4957.

8. Baum J, Richard D, Healer J, Rug M, Krnajski Z, Gilberger T-W, Green JL, Holder AA, Cowman AF. 2006. A conserved molecular motor drives cell invasion and gliding motility across malaria life cycle stages and other apicomplexan parasites. Journal of Biological Chemistry 281:5197–5208.

9. Ghosh AK, Coppens I, Gårdsvoll H, Ploug M, Jacobs-Lorena M. 2011. *Plasmodium* ookinetes coopt mammalian plasminogen to invade the mosquito midgut. Proceedings of the National Academy of Sciences 108:17153–17158.

10. Vega-Rodríguez J, Ghosh AK, Kanzok SM, Dinglasan RR, Wang S, Bongio NJ, Kalume DE, Miura K, Long CA, Pandey A, Jacobs-Lorena M. 2014. Multiple pathways for *Plasmodium* ookinete invasion of the mosquito midgut. Proceedings of the National Academy of Sciences 111:E492–E500.

11. Agrawal S, Chung D-WD, Ponts N, van Dooren GG, Prudhomme J, Brooks CF, Rodrigues EM, Tan JC, Ferdig MT, Striepen B, Le Roch KG. 2013. An apicoplast localized ubiquitylation system is required for the import of nuclear-encoded plastid proteins. PLoS Pathogens 9:e1003426–e1003426.

12. Yeh E, DeRisi JL. 2011. Chemical rescue of malaria parasites lacking an apicoplast defines organelle function in blood-stage *Plasmodium falciparum*. PLoS Biology 9:e1001138–e1001138.

13. Ganesan SM, Morrisey JM, Ke H, Painter HJ, Laroiya K, Phillips MA, Rathod PK, Mather MW, Vaidya AB. 2011. Yeast dihydroorotate dehydrogenase as a new selectable marker for *Plasmodium falciparum* transfection. Molecular and Biochemical Parasitology 177:29–34.

14. Storm J, Sethia S, Blackburn GJ, Chokkathukalam A, Watson DG, Breitling R, Coombs GH, Müller S. 2014. Phosphoenolpyruvate carboxylase identified as a key enzyme in erythrocytic *Plasmodium falciparum* carbon metabolism. PLoS Pathogens 10:e1003876–e1003876.

15. Jhun H, Walters MS, Prigge ST. 2018. Using lipoamidase as a novel probe to interrogate the importance of lipoylation in *Plasmodium falciparum*. mBio 9:e01872–18.

16. Collins CR, Das S, Wong EH, Andenmatten N, Stallmach R, Hackett F, Herman J-P, Müller S, Meissner M, Blackman MJ. 2013. Robust inducible Cre recombinase activity in the human malaria parasite *Plasmodium falciparum* enables efficient gene deletion within a single asexual erythrocytic growth cycle. Molecular Microbiology 88:687–701.

17. Knuepfer E, Napiorkowska M, van Ooij C, Holder AA. 2017. Generating conditional gene knockouts in *Plasmodium*-a toolkit to produce stable DiCre recombinase-expressing parasite lines using CRISPR/Cas9. Scientific Reports 7:3881–3881.

18. Birnbaum J, Flemming S, Reichard N, Soares AB, Mesén-Ramírez P, Jonscher E, Bergmann B, Spielmann T. 2017. A genetic system to study *Plasmodium falciparum* protein function. Nature Methods 14:450–456.

19. Armstrong CM, Goldberg DE. 2007. An FKBP destabilization domain modulates protein levels in *Plasmodium falciparum*. Nature Methods 4:1007–1009.

20. de Azevedo MF, Gilson PR, Gabriel HB, Simões RF, Angrisano F, Baum J, Crabb BS, Wunderlich G. 2012. Systematic analysis of FKBP inducible degradation domain tagging strategies for the human malaria parasite *Plasmodium falciparum*. PLoS One 7:e40981.

21. Muralidharan V, Oksman A, Iwamoto M, Wandless TJ, Goldberg DE. 2011. Asparagine repeat function in a *Plasmodium falciparum* protein assessed via a regulatable fluorescent affinity tag. Proceedings of the National Academy of Sciences 108:4411–4416.

22. Roberts AD, Nair SC, Guerra AJ, Prigge ST. 2019. Development of a conditional localization approach to control apicoplast protein trafficking in malaria parasites. Traffic 20:571–582.

23. Prommana P, Uthaipibull C, Wongsombat C, Kamchonwongpaisan S, Yuthavong Y, Knuepfer E, Holder AA, Shaw PJ. 2013. Inducible knockdown of *Plasmodium* gene expression using the *glmS* ribozyme. PloS One 8:e73783–e73783.

24. Ganesan SM, Falla A, Goldfless SJ, Nasamu AS, Niles JC. 2016. Synthetic RNA-protein modules integrated with native translation mechanisms to control gene expression in malaria parasites. Nature Communications 7:10727–10727.

25. Belmont BJ, Niles JC. 2010. Engineering a direct and inducible protein-RNA interaction to regulate RNA biology. ACS Chemical Biology 5:851–861.

26. Mair GR, Braks JAM, Garver LS, Wiegant JCAG, Hall N, Dirks RW, Khan SM, Dimopoulos G, Janse CJ, Waters AP. 2006. Regulation of sexual development of *Plasmodium* by translational repression. Science 313:667–669.

27. Amberg-Johnson K, Hari SB, Ganesan SM, Lorenzi HA, Sauer RT, Niles JC, Yeh E. 2017. Small molecule inhibition of apicomplexan FtsH1 disrupts plastid biogenesis in human pathogens. eLife 6:e29865.

28. Nasamu AS, Glushakova S, Russo I, Vaupel B, Oksman A, Kim AS, Fremont DH, Tolia N, Beck JR, Meyers MJ, Niles JC, Zimmerberg J, Goldberg DE. 2017. Plasmepsins IX and X are essential and druggable mediators of malaria parasite egress and invasion. Science 358:518–522.

29. Spillman NJ, Beck JR, Ganesan SM, Niles JC, Goldberg DE. 2017. The chaperonin TRiC forms an oligomeric complex in the malaria parasite cytosol. Cellular Microbiology 19:10.1111/cmi.12719.

30. Boucher MJ, Ghosh S, Zhang L, Lal A, Jang SW, Ju A, Zhang S, Wang X, Ralph SA, Zou J, Elias JE, Yeh E. 2018. Integrative proteomics and bioinformatic prediction enable a high-confidence apicoplast proteome in malaria parasites. PLoS Biology 16:e2005895–e2005895.

31. Garten M, Nasamu AS, Niles JC, Zimmerberg J, Goldberg DE, Beck JR. 2018. EXP2 is a nutrient-permeable channel in the vacuolar membrane of *Plasmodium* and is essential for protein export via PTEX. Nature Microbiology 3:1090–1098.

32. Ke H, Dass S, Morrisey JM, Mather MW, Vaidya AB. 2018. The mitochondrial ribosomal protein L13 is critical for the structural and functional integrity of the mitochondrion in *Plasmodium falciparum*. Journal of Biological Chemistry.

33. Walczak M, Ganesan SM, Niles JC, Yeh E. 2018. ATG8 is essential specifically for an autophagy-independent function in apicoplast biogenesis in blood-stage malaria parasites. mBio 9:e02021–17.

34. Florentin A, Stephens DR, Brooks CF, Baptista RP, Muralidharan V. 2019. The Clp system in malaria parasites degrades essential substrates to regulate plastid biogenesis. bioRxiv doi: 10.1101/718452:718452.

35. Istvan ES, Das S, Bhatnagar S, Beck JR, Owen E, Llinas M, Ganesan SM, Niles JC, Winzeler E, Vaidya AB, Goldberg DE. 2019. *Plasmodium* Niemann-Pick type C1-related protein is a druggable target required for parasite membrane homeostasis. eLife 8:e40529.

36. Rudlaff RM, Kraemer S, Streva VA, Dvorin JD. 2019. An essential contractile ring protein controls cell division in *Plasmodium falciparum*. Nature Communications 10:2181–2181.

37. Tang Y, Meister TR, Walczak M, Pulkoski-Gross MJ, Hari SB, Sauer RT, Amberg-Johnson K, Yeh E. 2019. A mutagenesis screen for essential plastid biogenesis genes in human malaria parasites. PLOS Biology 17:e3000136.

38. Fierro MA, Asady B, Brooks CF, Cobb DW, Villegas A, Moreno SNJ, Muralidharan V. 2020. An endoplasmic reticulum CREC family protein regulates the egress proteolytic cascade in malaria parasites. mBio 11:e03078–19.

39. Gupta A, Bokhari AAB, Pillai AD, Crater AK, Gezelle J, Saggu G, Nasamu AS, Ganesan SM, Niles JC, Desai SA. 2020. Complex nutrient channel phenotypes despite Mendelian inheritance in a *Plasmodium falciparum* genetic cross. PLoS Pathogens 16:e1008363–e1008363.

40. Ling L, Mulaka M, Munro J, Dass S, Mather MW, Riscoe MK, Llinás M, Zhou J, Ke H. 2020. Genetic ablation of the mitoribosome in the malaria parasite *Plasmodium falciparum* sensitizes it to antimalarials that target mitochondrial functions. Journal of Biological Chemistry doi: 10.1074/jbc.RA120.012646

41. Nessel T, Beck JM, Rayatpisheh S, Jami-Alahmadi Y, Wohlschlegel JA, Goldberg DE, Beck JR. 2020. EXP1 is required for organisation of EXP2 in the intraerythrocytic malaria parasite vacuole. Cellular Microbiology 22:e13168.

42. Polino AJ, Nasamu AS, Niles JC, Goldberg DE. 2020. Assessment of biological role and insight into druggability of the *Plasmodium falciparum* protease Plasmepsin V. ACS Infectious Diseases 6:738–746.

43. Zhang M, Wang C, Otto TD, Oberstaller J, Liao X, Adapa SR, Udenze K, Bronner IF, Casandra D, Mayho M, Brown J, Li S, Swanson J, Rayner JC, Jiang RHY, Adams JH. 2018. Uncovering the essential genes of the human malaria parasite *Plasmodium falciparum* by saturation mutagenesis. Science 360:eaap7847.

44. Spork S, Hiss JA, Mandel K, Sommer M, Kooij TWA, Chu T, Schneider G, Maier UG, Przyborski JM. 2009. An unusual ERAD-like complex is targeted to the apicoplast of *Plasmodium falciparum*. Eukaryotic Cell 8:1134–1145.

45. Adjalley SH, Johnston GL, Li T, Eastman RT, Ekland EH, Eappen AG, Richman A, Sim BKL, Lee MCS, Hoffman SL, Fidock DA. 2011. Quantitative assessment of Plasmodium falciparum sexual development reveals potent transmission-blocking activity by methylene blue. Proceedings of the National Academy of Sciences 108:E1214.

46. Ghorbal M, Gorman M, Macpherson CR, Martins RM, Scherf A, Lopez-Rubio JJ. 2014. Genome editing in the human malaria parasite *Plasmodium falciparum* using the CRISPR-Cas9 system. Nature Biotechnology 32:819–821.

47. Swift RP, Rajaram K, Liu HB, Dziedzic A, Jedlicka AE, Roberts AD, Matthews KA, Jhun H, Bumpus NN, Tewari SG, Wallqvist A, Prigge ST. 2020. A mevalonate bypass system facilitates elucidation of plastid biology in malaria parasites. PLoS Pathogens 16:e1008316–e1008316.

48. Deitsch K, Driskill C, Wellems T. 2001. Transformation of malaria parasites by the spontaneous uptake and expression of DNA from human erythrocytes. Nucleic Acids Research 29:850–853.

